# Accurate label-free quantification by directLFQ to compare unlimited numbers of proteomes

**DOI:** 10.1101/2023.02.17.528962

**Authors:** Constantin Ammar, Julia Patricia Schessner, Sander Willems, André C. Michaelis, Matthias Mann

## Abstract

Recent advances in mass spectrometry (MS)-based proteomics enable the acquisition of increasingly large datasets within relatively short times, which exposes bottlenecks in the bioinformatics pipeline. Whereas peptide identification is already scalable, most label-free quantification (LFQ) algorithms scale quadratic or cubic with the sample numbers, which may even preclude the analysis of large-scale data. Here we introduce directLFQ, a ratio-based approach for sample normalization and the calculation of protein intensities. It estimates quantities via aligning samples and ion traces by shifting them on top of each other in logarithmic space. Importantly, directLFQ scales linearly with the number of samples, allowing analyses of large studies to finish in minutes instead of days or months. We quantify 10,000 proteomes in 10 minutes and 100,000 proteomes in less than two hours - thousand-fold faster than some implementations of the popular LFQ algorithm MaxLFQ. In-depth characterization of directLFQ reveals excellent normalization properties and benchmark results, comparing favorably to MaxLFQ for both data-dependent acquisition (DDA) and data-independent acquisition (DIA). Additionally, directLFQ provides normalized peptide intensity estimates for peptide-level comparisons. It is available as an open-source Python package and as a GUI with a one-click installer and can be used in the AlphaPept ecosystem as well as downstream of most common computational proteomics pipelines.

---

Mass Spectrometry (MS)-based proteomics is the method of choice for global analysis of the proteome^1^, including applications in clinical^2,3^, single cell^4,5^ and spatial^6^ proteomics. Modern MS-proteomics workflows are increasingly quantitative, meaning that the major insights derived from the experiments base on observed changes in protein abundances^7^. Appropriate computational processing is therefore key for obtaining unbiased quantitative protein values from the raw ion intensities acquired by the instrument^8,9^. Conceptually, the computational processing of MS proteomics data is comprised of identification and quantification. In the identification step, the ion signals acquired by the MS instrument are assigned to their most probable peptides and proteins using statistical models. In the quantification step, the intensities of the ion signals are used to derive meaningful proxies for protein (or peptide) abundances.

In this study, we focus on the quantification step, which again consists of two essential parts: the normalization of systematic biases between samples and the generation of peptide and protein intensities from the underlying ion intensities. A variety of methods have been established for quantification^10–14^, of which the MaxLFQ approach^10^ is one of the most widely used and implemented in most of the current computational proteomics pipelines^15–20^. One reason for the popularity of MaxLFQ is that it addresses major pitfalls in proteomics quantification by accounting for the fact that different peptides belonging to the same protein can have very different base intensities, for example due to differing ionization efficiencies^21^. Additionally, it is robust against missing values for normalization (differing sample depths) as well as in protein intensity estimation. MaxLFQ achieves this by solving equation systems for both steps^10^. However, the number of terms and equations scales quadratically with the number of samples, leading to challenges in execution time as well as high overall complexity of the approach. Although this can be alleviated by faster implementations of the algorithms for quadratic optimization^17,22^, the issue of quadratic increase remains, imposing an upper limit for feasible sample numbers.

To mitigate these issues, we introduce directLFQ as a simple and direct method for sample normalization and protein intensity estimation. We re-frame the normalization problem by using the concept of “intensity traces”, which are aligned by a single scaling factor per intensity trace. By re-framing the problem in such a way, we reduce the quadratic to a linear scaling (i.e. 10 times more samples only take 10 times longer execution time), for any number of samples. Comparing directLFQ to MaxLFQ in several quantitative benchmarking sets, we show an overall better performance of directLFQ, for both data-dependent acquisition (DDA) and data-independent-acquisition (DIA) datasets. In a complex spatial proteomics dataset, we observe that several aspects of the biology are better resolved with directLFQ.

The code of directLFQ is openly available on GitHub (https://github.com/MannLabs/directlfq), with easy access for all types of users (graphical user interface (GUI) for end users, as well as a command line interface (CLI) and a Python application programming interface (API)). Supported input formats include AlphaPept^17^, MaxQuant^16^, Spectronaut^18^, DIA-NN^19^ and IonQuant/FragPipe^20^. The underlying algorithm allows for easy adaptation to other software pipelines.

## EXPERIMENTAL PROCEDURES

### directLFQ analysis of MaxQuant files

As examples of Data-Dependent Acquisition (DDA), MaxQuant result files were (re)processed in this study by calling directLFQ on the evidence.txt output file. The proteinGroups.txt file was used as an additional input for protein group mapping.

### Dynamic Organellar Maps analysis

For analysis of the DOM data, the original MaxQuant output files from the study of Schessner et al.^23^ acquired on an Orbitrap Exploris 480 mass spectrometer were used. directLFQ was called on the MaxQuant files as described above. Subsequently, in order to fulfill the formatting requirements of the DOM-QC analysis tool, an adapted directLFQ file was created, where columns from the proteinGroups.txt file were added to the directLFQ file. The adapted directLFQ file as well as the original proteinGroups.txt file were both uploaded to the DOM-QC tool (https://domqc.bornerlab.org/QCtool) and a .yaml file containing the aligned comparison results was downloaded. Plots were created on a local machine based on the DOM-QC code available on GitHub (https://github.com/valbrecht/SpatialProteomicsQC).

### DDA mixed species benchmark

For the DDA mixed species benchmark described below, the MaxQuant output files of the “HeLa - *E. coli*” dataset acquired by our group (Meier et al.^24^) on a Q Exactive HF mass spectrometer were downloaded from the respective PRIDE^25^ repository PXD006109. directLFQ was called on the MaxQuant files as described above. For benchmarking, a subset of six files was used that had been acquired in standard DDA mode (not in BoxCar mode). Three files contained six times the amount of *E. coli* as compared to the other three. For analysis, the median log2 transformed LFQ intensity of the high *E. coli* set was obtained and subtracted from the median log2 transformed LFQ intensity of the low *E. coli* set. This resulted in one log2 fold change per protein. For analysis, these fold changes were plotted as violin plots or scattered against the mean of the log2 transformed LFQ intensities over all samples.

### Consistency analysis of 200 HeLa files

To assess the performance of directLFQ on larger datasets, the MaxQuant output files of the “200 HeLa” dataset by Bian et al.^26^ acquired on a Q Exactive HF-X mass spectrometer was downloaded from the corresponding PRIDE repository PXD015087. directLFQ was called on the MaxQuant files as described above. The directLFQ output file and the proteinGroups.txt file were used for further analysis. The CV was calculated for every protein over the 200 HeLa files and the resulting distributions of CVs were compared.

### DIA mixed species benchmark

For the DIA mixed species benchmark, six .raw files corresponding to the “large-FC” dataset by Huang et al.^27^ acquired on a Q Exactive HF mass spectrometer were downloaded from the PRIDE repository PXD016647. The raw files were analyzed using Spectronaut 15 in directDIA mode based on the Uniprot databases UP000000625, UP000005640, UP000002311 and UP000001940.

The Spectronaut report was exported in the format specified in the spectronaut_tableconfig_fragion.rs file available on the directLFQ GitHub repository and accessible via the directLFQ GUI. directLFQ was called on the exported Spectronaut results file and the protein intensities were estimated based on a combination of the MS1 isotope intensities as well as the fragment ion intensities (i.e. each fragment ion and MS1 isotope corresponded to one independent intensity trace). The iq package was called on the same report file using the “iq::process_long_format” command.

The raw files mapped to two conditions S1 and S2, with three samples in S1 and three in S2. *S. cerevisiae* and *C. elegans* proteins had different abundances in S1 and S2 (ratios 2 and 0.77, respectively), while *H. sapiens* remained constant. As described above for the DDA data, the median log2 transformed intensity of each protein was obtained and these intensities were subtracted between S1 and S2. The intensity estimate for each protein was the mean log2 transformed intensity over all conditions.

### Consistency analysis of clinical DIA study

The DIA-NN results file of the study of Demichev et al.^28^ (acquired on a TripleTOF 6600 mass spectrometer) was downloaded from the PRIDE repository PXD029009. directLFQ was called on the output file, using the “MS1.Area” as well as the “Fragment.Quant.Raw” columns for creating ion intensities. Each fragment ion as well as the MS1.Area corresponded to one independent intensity trace. The DIA-NN implemented MaxLFQ results were obtained via the “Genes.MaxLFQ” column. The iq package was called on the appropriately reformatted report using the “iq::process long format” command.

The dataset contained several types of quality control (QC) samples. As a quality measure the CV for each type of quality control sample was obtained, for every protein. The distributions of all CVs together were then used for comparing the different LFQ approaches.

### Normalization analysis

To test normalization on a challenging dataset, the MaxQuant result files of the tissue dataset by Wang et al.^29^ acquired on an Orbitrap Fusion Lumos mass spectrometer were downloaded from the PRIDE repository PXD010154. Precursor (sequence and charge) intensities were extracted from the evidence.txt file. directLFQ normalization was performed by calling the directLFQ normalization class on the precursor table. Median normalization was performed using R code for normalization provided together with the iq package (https://cran.r-project.org/web/packages/iq/vignettes/iq.html).

The dataset consisted of several different tissue measurements, from which “lung” was chosen as the reference tissue. The precursor intensities were log2 transformed and the precursor intensity of the lung tissue was subtracted from the precursor intensities of the other conditions. This resulted in a distribution of ratios (one ratio per precursor) relative to lung for each tissue. The boxplots in the main text below contain the set of precursors that do not have any missing values, while normalization was performed on the complete dataset. We chose the precursors with no missing values for visualization because they are more likely to reflect the true abundance changes between the tissues.

### Execution time analysis

Execution time comparison was performed on a medium-strength computer cluster (128 GB RAM, 64 logical processors @2.4GHz). Execution times of up to 10,000 samples on this cluster were comparable to execution times on a state-of-the-art MacBook (MacBook Pro 16-inch, 2021, M1 Max, 64GB RAM). The MaxLFQ reference runs were performed in the scope of an interactomics study^30^ on a comparable cluster (512 GB RAM, 40 logical processors @2.2GHz). directLFQ runs on CPUs with multiprocessing enabled per default and the option to manually set the number of cores to use.

Different sample sizes were simulated by using a template dataset which contained N samples. To simulate M>N samples, we duplicated the N samples as often as was necessary to reach M. In the case of M<N, we left out as many samples as necessary, by taking the first M columns in the template.

The template dataset for DDA data was based on the interactomics study and the template dataset for DIA data was based on an example dataset provided together with the iq package (https://github.com/tvpham/iq/releases/download/v1.1/Bruderer15-DIA-longformat-compact.txt.gz).

We called directLFQ on the templates using the “lfq_benchmark.LFQTimer” class. For iq, we used the system.time command on the “iq::fast_preprocess” and the “iq::fast_MaxLFQ” commands. The data was adjusted to adhere to the format necessary for the “iq::fast_preprocess” command.

For the 100,000 sample datasets, the memory limit of 128GB was surpassed, which is why we decreased the number of proteins in the template file by a factor of 4 and multiplied the resulting execution time by four. As the proteins are processed independently of each other this should be a realistic estimation of the true execution time.

## RESULTS

### Reframing the normalization problem with intensity traces

The underlying idea of directLFQ approach is to frame the set of samples or ions to be normalized as a set of intensity traces. In the case of sample normalization, such a trace consists of all ion intensities measured in a sample (**Figure 1A**). Each point in the trace has the coordinates: (ion identifier, log2(intensity) of ion). For example, the trace representing sample 1 might be precursors VTTHPLAK_CHARGE2 with log2(intensity) of 23, VTVAGLAGK_CHARGE3 with log2(intensity) of 25 and so on. In sample 2, these same peptides will have differing intensities, resulting in a slightly different trace. There are as many traces as there are samples in the dataset.

**Figure 1:**
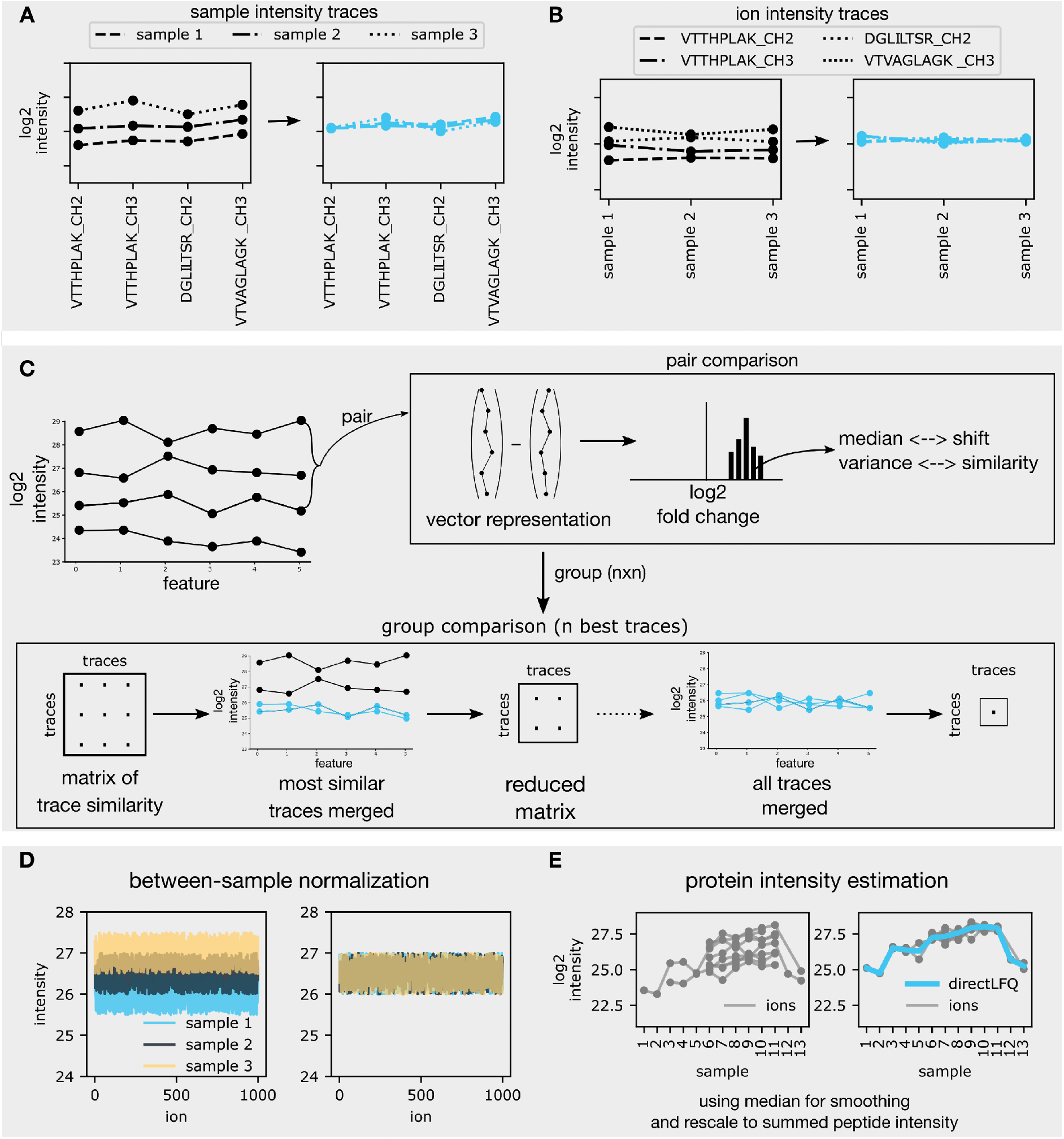
The directLFQ approach. Objects to be normalized are intensity traces that can be shifted. A) Between-sample normalization, where each trace represents a sample and each element of the trace is a peptide’s log2 intensity. Traces are shifted on top of each other (blue) as described below. B) Protein intensity estimation, where each trace belongs to a peptide and each element of the trace is a sample’s log2 intensity. C) The shifting process. Traces are compared in a pairwise fashion by subtracting the intensities and extracting the median and variance of the resulting difference distribution (top). The most similar samples are shifted (indicated in blue) and a merged sample is created. The process is repeated on a now smaller similarity matrix until only one trace remains (bottom). A more realistic example for between-sample normalization is given in D) and for subsequent protein intensity estimation in E).

In the case of protein intensity estimation, each trace represents specific precursor ions (DDA) or also the transitions (DIA), and each trace contains all intensities of these ions over the different samples (**Figure 1B**). For example, the protein containing the precursors VTTHPLAK_CHARGE2, VTTHPLAK_CHARGE3, DGLILTSR_CHARGE2 and VTVAGLAGK _CHARGE3, would have four ion intensity traces with each point in a trace defined by the coordinates: (sample number, log2(intensity)).

Note that the concept of intensity traces requires log transformation, because the shape of the intensity trace is then determined by the relative changes between samples, independent of base intensity. For simplicity, we denote the x-axis components of the intensity traces as *features*, which could be either samples, precursor ions, or transitions, depending on the context.

### Shifting intensity traces to compare relative changes

For the directLFQ approach, we frame sample normalization and protein intensity estimation as problems of intensity traces that need to be shifted (**Figure 1C**). We do not alter the shape of each intensity trace, but instead make the shapes of the intensity traces comparable to each other. This is achieved by adding one scaling factor to each intensity trace (corresponding to multiplication because of the log transformation). The scaling factors are chosen to minimize the distance of the intensity traces from each other (see section below). This way the shapes of the intensity traces are preserved, but the systematic shifts between the traces are corrected for. In the case of ‘between sample normalization’ this means that systematic intensity shifts between samples are corrected without changing the relative intensities of ions within each sample. In the case of ‘protein intensity estimation’ this means that systematic biases between ions - due to differing ionization efficiencies of precursors in case of protein intensity estimation - are corrected for, again preserving the relative abundance changes of the ions between samples.

### A systematic approach to shifting intensity traces

As mentioned, the core concept of directLFQ is to shift intensity traces on top of each other. A wide variety of methods can implement this shifting. Most simply, one original or an average/median trace could be selected and all traces could be shifted towards that trace. Alternatively, one could treat this as a minimization problem e.g., using a quadratic solver. Based on our previous work with the MS-EmpiRe^31^ package, we here chose to use pair-wise comparisons for shifting using an adapted single linkage approach. This means that we compare all pairs of traces to each other as follows: Each of the n traces is considered a vector of log2(intensities) with as many elements as features (**Figure 1C**, upper right). In this vector, missing values are encoded as NaN (Not a Number). When subtracting both vectors, this results in a distribution of fold changes. From this distribution we then extract the median and the variance. The median estimates any systematic shift between the intensity traces, whereas the statistical variance reflects the overall divergence of the traces. We shift the whole set of traces using an iterative procedure as shown at the bottom of **Figure 1C**: Given n traces, we first collect two *n* · *n* (half-) matrices, one containing the variances of all pairs of intensity traces and one containing the medians. From the variance matrix, we extract the pair of intensity traces that is most similar - the smallest element in the matrix. One of the intensity traces in the pair is rescaled to the other by adding the corresponding median shift from the median half matrix. After scaling, the pair of traces is combined, creating a new and more stable ‘averaged trace’ that replaces the pair.

After also recalculating the affected elements of the median and the variance matrices, the procedure is repeated on the resulting (*n* – 1) · (*n* – 1) matrices until all *n* intensity traces are merged. Each time an averaged trace is shifted, the corresponding scaling factors need to be propagated to all samples underlying the averaged trace. To this end, the scaling factors are tracked throughout the procedure and then used to shift the original intensity traces on top of each other.

With this approach, the most similar samples are merged first, decreasing the possible error in each shift. Creating the average intensity trace in this way has two conceptual benefits: first, it should be more stable than the single-intensity traces, because it is an average of multiple traces. Secondly, creating the average trace mitigates the missing value problem. If an intensity at a certain position is missing in one vector, but present in another, the averaged vector will automatically be filled with the intensity that is not missing.

As this comparison scales quadratically with *n*, we define upper limits *n_max_* (*n_max_* = 50 for sample normalization and *n_max_* = 10 for protein intensity estimation). If there are more than *n_max_* intensity traces, we construct an average from the *n_max_* intensity traces with the lowest numbers of missing values and shift all remaining intensity traces towards this average. The reasoning behind this is that an average constructed from *n_max_* traces should be sufficiently stable and complete to enable precise sample shifting. We tested this assumption for a case of sample normalization, where the number of samples (1600) is much larger than *n_max_*. This showed that the variance between samples did not depend on *n_max_*, from 5 to 200, indicating that the most complete traces picked first by the algorithms are already an excellent basis for robust normalization (**Supplementary Figure 1**). After having created the average, the further shifting steps are trivial and computationally inexpensive. We name the combination of the ion trace concept and the trace shifting algorithm ‘directLFQ’, because it addresses possible quantification biases most directly.

### Between-sample normalization

With the directLFQ algorithm in hand, we perform between-sample normalization as the first step of the quantification workflow. Here each sample is rescaled as described above and visualized in **Figure 1D**, adjusting for biases such as due to differences in sample loading or different performance of the mass spectrometer. The underlying assumption is that the majority of proteins are not regulated between samples, because we use the median between intensity traces as a scaling factor. This is a common assumption based on biological observations of proteome regulation and implicit in many normalization algorithms, including MaxLFQ^10^. In case the majority of proteins is regulated, instead of the median, one could use the mode as the metric (the most often observed change between samples). However, this is more challenging to estimate in noisy data and is therefore often less stable than the median. The most common biological reason that our assumption of an unchanging majority of proteins does not hold, would be that a particular group of ‘uninteresting’ proteins such as contaminants or extracellular matrix, for instance, are present in a subset of samples. These can then be excluded from the intensity traces in directLFQ by providing a subset of ‘housekeeping proteins’^32,33^ to perform normalization on. directLFQ offers the option to pass a set of such proteins and normalization will be performed using this subset.

### Protein intensity estimation

After sample normalization, directLFQ estimates the most likely protein profile as illustrated in **Figure 1E**. After the ion intensity traces are shifted on top of each other, the median intensity of each sample is an estimate for the relative protein intensity. The protein intensity profile is then transformed back from log2 transformed intensities into linear space and multiplied by a single factor representing the overall protein abundance. This is the sum of all linear ion intensities over all samples for the given protein divided by the sum of all linear protein intensities over all samples. In this way, the overall peptide intensity is retained, as in the MaxLFQ algorithm^10^.

### Handling DDA, DIA and other types of acquisition data

We have tested and benchmarked directLFQ on DDA as well as DIA data, with DDA having quantitative data only on the MS1 level and DIA at the MS1 and MS2 levels. For DDA data processing, we build ion intensity traces based on the MS1 intensities of the charged precursors. For DIA data, we build intensity traces based on the MS1 intensities of the charged precursors as well as on the MS2 level fragment ion intensities. Thus DIA multiplies the number of data points available for quantification, which we find to stabilize the protein intensity estimation (**Supplementary Figure 2**). The directLFQ algorithm can also readily be applied to different types of quantitative proteomics, including isotope label-based methods such as Tandem Mass Tags^34^ (TMT). For TMT experiments contained in a single plex, the algorithm can be applied. In case there are more conditions than TMT channels, the corresponding experiments should be ‘channel normalized’ before using directLFQ.

### Timing directLFQ on up to 100,000 samples

As described above, the directLFQ algorithm scales linearly with the number of samples, which should in principle allow to quantify arbitrarily large numbers of samples in a reasonable time. To test this in a controlled manner and in the absence of extremely large datasets, we used real datasets as templates and simulated increasing numbers of samples by replication (Experimental Methods). For any of the methods that we compared, we used the elapsed execution times for normalization and protein intensity estimation for benchmarking, without data loading and data output.

**Figure 2A** depicts the timing results for the very large dataset of our recent yeast interactome project^30^ as processed by MaxQuant. The option “fastLFQ”^10^ was enabled in processing with MaxLFQ. Its execution times increased quickly and reached two weeks of processing time at around 2,000 samples after which we broke off the computation. In contrast, directLFQ took around two minutes for the same number of samples. To keep execution times reasonable, we omitted larger samples with MaxLFQ and further scaled up the number of samples for directLFQ to 10,000 and 100,000 by replication, which took 10 minutes and 100 minutes respectively (Experimental Methods).

**Figure 2:**
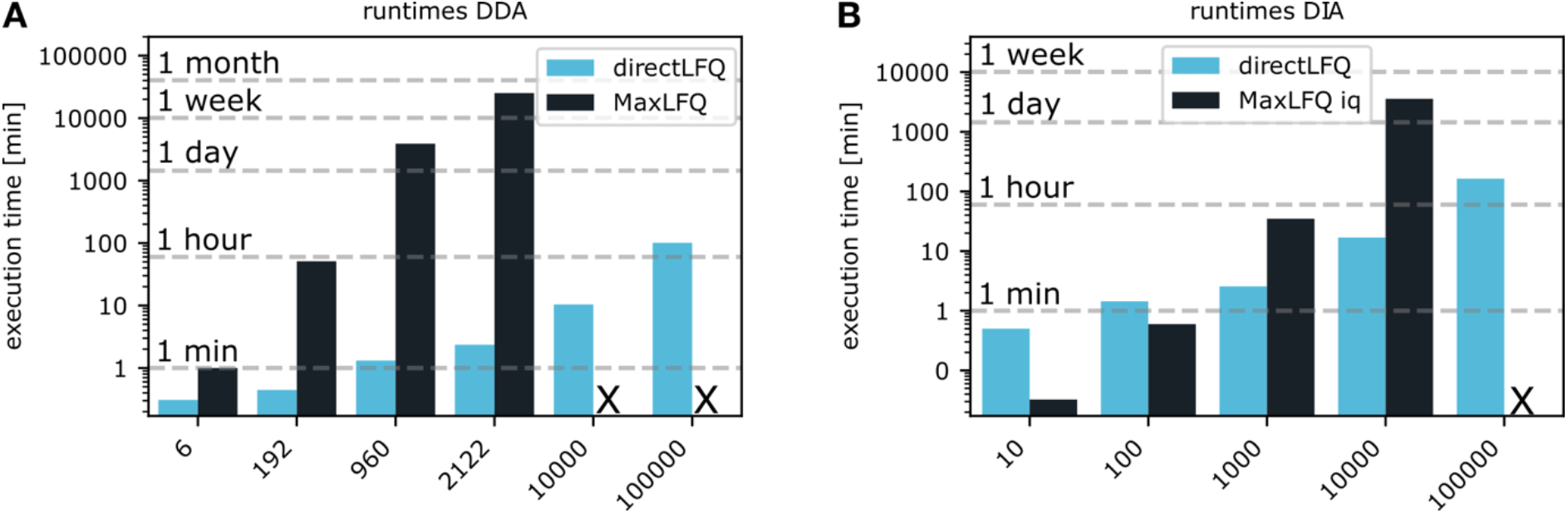
Processing times for different methods and sample sizes. directLFQ scales (sub-) linearly for A) DDA data and B) DIA data, resulting in more than 1000-fold faster execution times than MaxLFQ. X in the plot marks instances with prohibitive calculation times.

For DIA data, we instead compared against the fast C++ based implementation of the popular R package iq, as recommended for processing DIA data. Due to its very fast implementation of the quadratic LFQ algorithm, it took only 2s on its small 10 sample dataset, compared to 30s for directLFQ. While this difference is inconsequential in routine proteomics practice, we do observe the expected non-linear increase in execution time in function of the number of samples and 10,000 samples already require a processing time of 2.5 days, while directLFQ takes around 16 minutes. We further scaled directLFQ up to 100,000 samples which took only around 160 minutes. Note that directLFQ is only parallelized for the CPU, whereas implementation on GPUs would further drastically shrink processing time.

The increased processing time of directLFQ in DIA compared to DDA is due to the fact that we have one intensity trace for each fragment ion, increasing the total number of traces to be processed.

### Applying directLFQ to benchmarking datasets

Next, we benchmark directLFQ on several public datasets that are meant to directly assess quantification performance. We first applied directLFQ to a mixed-species (*Homo sapiens* (*H. sapiens*) and *Escherichia coli* (*E. coli*)) dataset acquired in DDA mode by our group^24^. A total of six samples were measured, containing identical amounts of *H. sapiens* cell lysate, while three of them had sixfold more *E. coli* than the others. Comparing the two groups with three samples each, a ratio around one would be expected for the *H. sapiens* proteins and a ratio of six for the *E. coli* proteins. The better the quantification is, the closer the proteins should be to the expected ratios. The directLFQ derived protein intensities align well around the expected ratios, with substantially reduced outliers and lower standard deviation (0.57 instead of 0.86 for *E. coli*) as compared to MaxLFQ Additionally, the *H. sapiens* proteins are centered around the expected ratio in directLFQ, while we see systematic deviation in MaxLFQ (**Figure 3A**).

**Figure 3:**
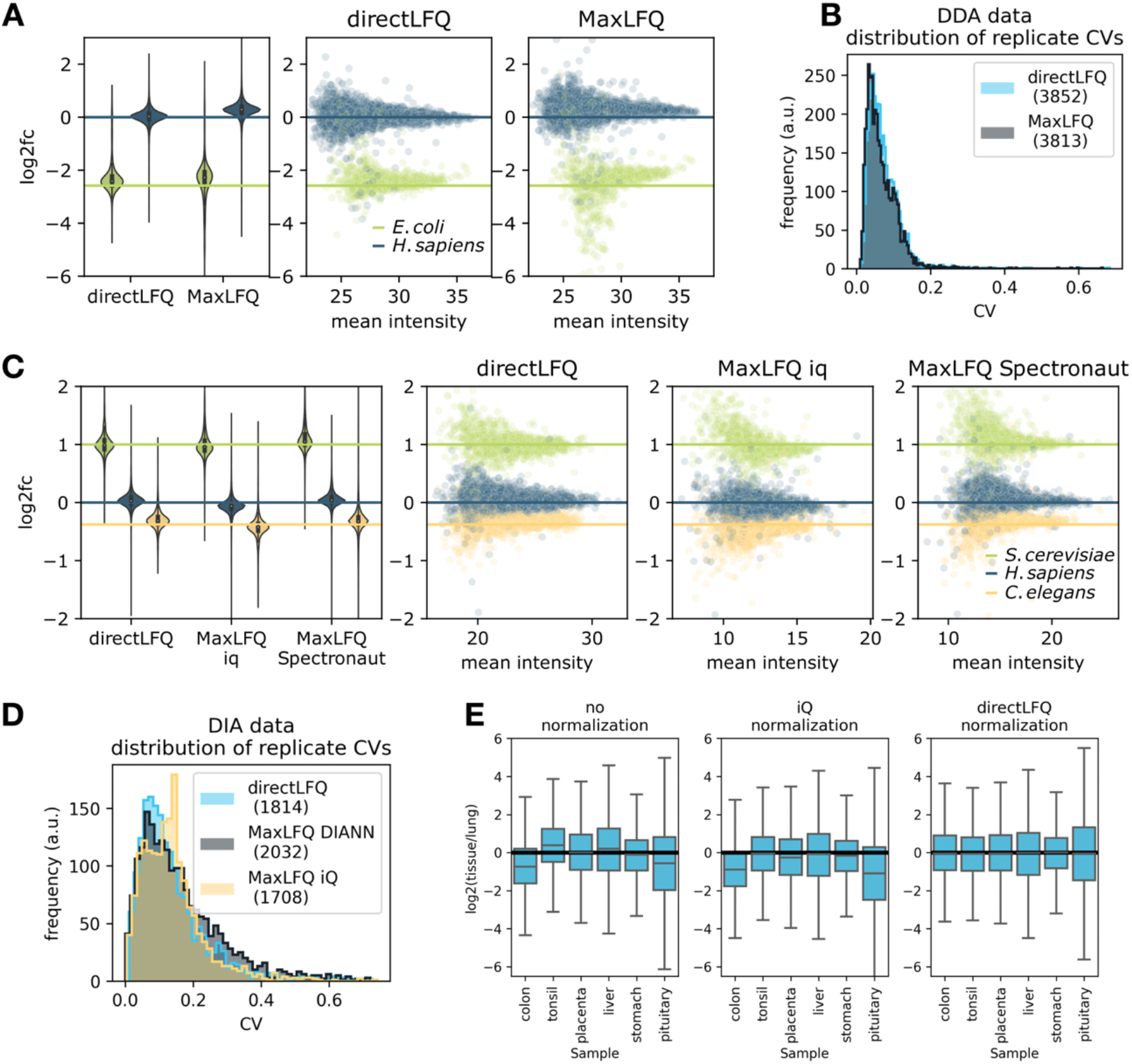
Applying directLFQ to different benchmarking datasets. A) Mixed-species DDA dataset processed with directLFQ and MaxLFQ. *E. coli* proteins should align along a log2 ratio of −2.59 (blue line), and *H. sapiens* proteins should align along a ratio of 0. B) Distribution of CV values on a DDA dataset with 200 replicate HeLa samples, with very similar results for directLFQ and MaxLFQ. C) Mixed-species DIA dataset processed with directLFQ and two MaxLFQ implementations (iq and Spectronaut). Expected log2 ratios for *S. cerevisiae, H. sapiens* and *C. elegans* proteins are −0.38, 0 and 1, respectively. D) Distribution of CV values between technical repeat samples from a ~900 sample clinical DIA dataset, processed with directLFQ and two MaxLFQ implementations (iq and DIA-NN) with comparable results for all approaches. E) Testing directLFQ precursor normalization on a challenging tissue dataset and comparing against standard median normalization. After normalization, all boxes should be aligned around 0, which is the case for directLFQ but not for the median normalization approach.

To test the performance of directLFQ on larger experiments, we applied it to a published 200 sample HeLa technical dataset^29^ (**Figure 3B**). As the samples should be identical and each protein should therefore not significantly change between samples, we used the coefficient of variation (CV) of each protein as a quality measure. Our results show CV distributions of directLFQ and MaxLFQ that are nearly identical.

To test directLFQ on DIA data, we applied it to a three-species dataset that was acquired by Huang et al.^27^ that we processed with Spectronaut^18^ (*S. cerevisiae, H. sapiens* and *C. elegans*) (Experimental Methods). It consists of six samples, split into two conditions, with expected ratios 0.77, 1 and 2 (yeast, human, C. *elegans*). We compared directLFQ against the above mentioned iq package and the MaxLFQ implementation of MaxLFQ implementation of Spectronaut. Both the expected ratios and the spread around them were better for directLFQ. Standard deviations for *S. cerevisiae* were 0.21, 0.26 and 0.31 for directLFQ, iq and Spectronaut respectively. iq showed a systematic offset for the *H. sapiens* proteins which is not visible for directLFQ and Spectronaut (**Figure 3C**).

We next tested performance on a clinical DIA dataset consisting of almost 1,000 Covid 19 plasma proteomes^28^, which had been processed with the software DIA-NN^19^. This included many quality control (QC) samples, which ideally should show no variability between runs, allowing us to use the CVs on the quality control samples as a quality measure of quantification. Comparing directLFQ against iq and DIA-NN, revealed that the distributions of CVs are in a similar range, with iq showing an anomalous peak at a CV of 0.18 and directLFQ and DIA-NN being comparable (**Figure 3D**).

Lastly, we tested the sample normalization algorithm of directLFQ on a tissue dataset. We chose tissues because they often have strong differences in their proteome composition and therefore constitute a challenging normalization benchmark. In a first step, we used a lung proteome as a reference and calculated the ratio to lung for each precursor in each tissue. This results in a set of ratios for each of the tissues, with systematic shifts between them (**Figure 3E).** The objective of a normalization function is then to assign constant scaling factors to each tissue, such that they are optimally aligned. Clearly, a simple median normalization, as for example implemented in the iq package, does not suffice to align the datasets (**Figure 3E**, middle), while directLFQ normalization results in well centered distributions (**Figure 3E**, right). This demonstrates that the directLFQ normalization strategy is effective in correcting systematic biases between samples. As noted above, such normalization is only recommended if less than half of the quantified proteins are substantially regulated. If this is not the case directLFQ provides the option to specify a protein subset to normalize on.

### Applying directLFQ to organellar maps data

To test directLFQ in a sophisticated cell biological situation involving proteomic separation at different time points, we analyzed a dynamic organellar maps (DOM) dataset from Schessner et al.^23^. In this dataset, cells were mechanically lysed and the resulting cellular compartments were separated using differential centrifugation^23^. Different *fractions* of the separated sample were measured with DDA. In these organellar maps, proteins are most abundant in the fractions that correspond to their localization (for instance, Golgi apparatus). Furthermore, proteins belonging to the same protein complex or organelle are expected to have similar intensity profiles. This DOM dataset is a good benchmark for the performance of the directLFQ algorithm because of this complex biological dataset with many distinct outcomes. Additionally, the paper provides the DOM-QC benchmarking tool that is meant to assess quantification performance on this type of data, which we use in the evaluations shown.

Principal-component-analysis (PCA) by DOM-QC of the directLFQ processed data visibly separates different organellar parts of the cell (**Figure 4A**). Comparing the directLFQ results to the MaxLFQ results originally used in the publication suggests a more consistent clustering of the directLFQ data. This is especially visible for the endosome, peroxisome or Golgi proteins. The overall spread of the MaxLFQ data is higher, which could either have biological or algorithmic reasons. However, investigating the proteins at the extremes of the PCA indicates that directLFQ accurately reflects the changes visible on precursor level, while MaxLFQ appears to over-estimates protein ratios (**Supplementary Figure 3**).

**Figure 4:**
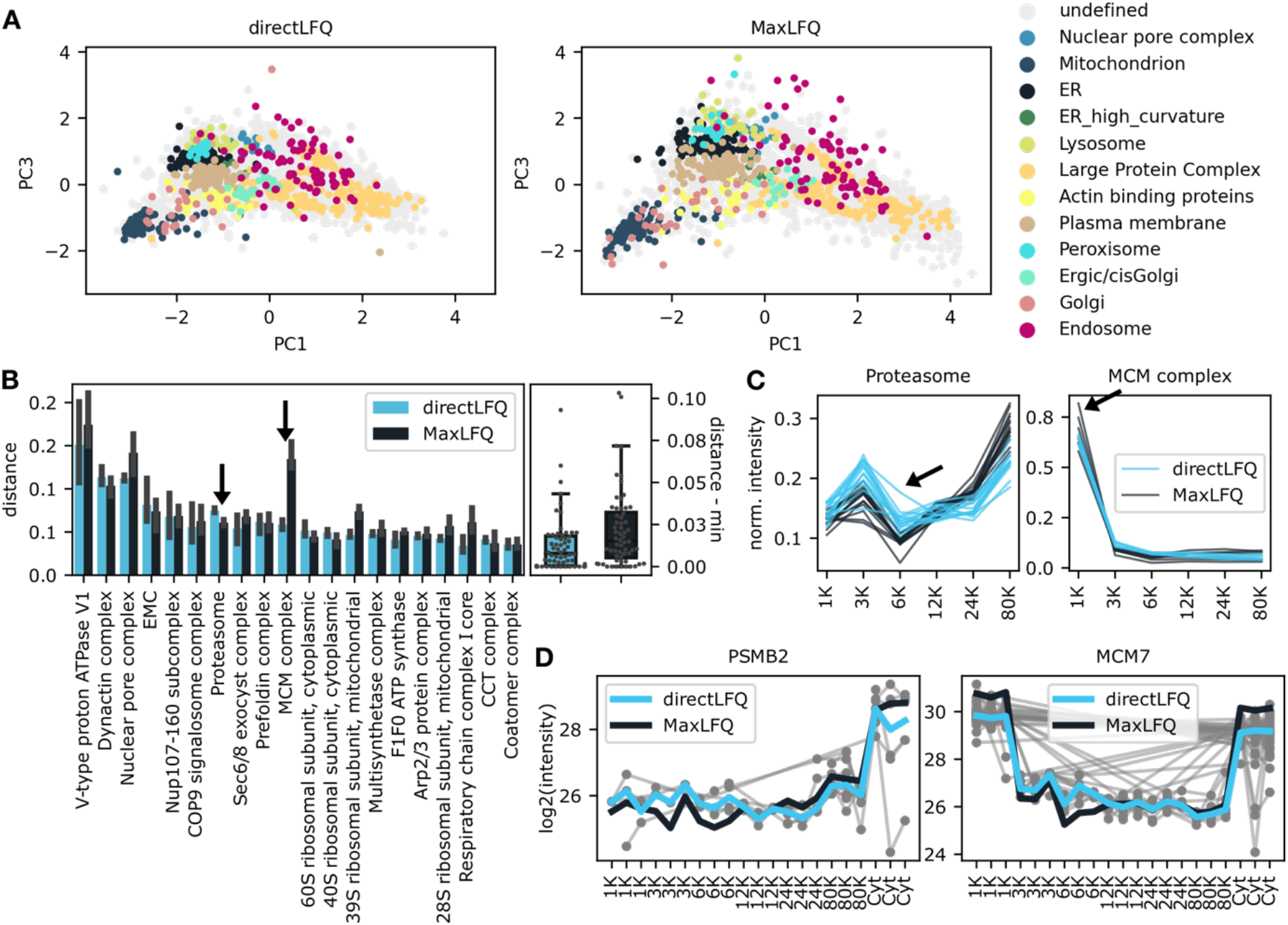
Applying directLFQ to dynamic organellar maps (DOM) data from ref.^23^ and comparing to MaxLFQ. A) PCA maps of the DOM data in which protein clusters are color coded. Several clusters such as Golgi and Mitochondrion are separated more clearly with directLFQ. B) Quantitative assessment of the similarity of the intensity profiles of protein clusters (lower distance means better consistency). On the left, the distances with error bars are displayed for each tested protein cluster. The arrows indicate the two clusters MCM complex and Proteasome where directLFQ and MaxLFQ perform best, respectively. On the right, the normalized distances are compared to each other as boxplots. C) Protein intensity profiles of these two clusters. One outlier trace in each cluster is marked by an arrow. D) Visualization of the protein profiles over all replicates together with the underlying ion data. The traces show that directLFQ faithfully represents the underlying data.

To further explore the differences in protein cluster consistencies, we used the DOM-QC tool to quantify the consistency of protein profiles for proteins belonging to the same complex (**Figure 4B**). On the default list of complexes provided by DOM-QC, the overall distance within clusters of directLFQ is consistently better (lower values) with few exceptions.

The protein intensity profiles of the MCP complex had the best results in directLFQ compared to MaxLFQ and those of the proteasome complex were best in MaxLFQ (**Figure 4C** and indicated by arrows in **Figure 4B**). In the MCM complex, where directLFQ compared favorably, we see that the traces of directLFQ are more tightly aligned, with almost identical profiles. For the Proteasome, where MaxLFQ compared favorably, we see that directLFQ has a deviating trace. Zooming in further, we examined the most deviating protein MCM7 of the MCM complex in the MaxLFQ data. This indicated that the algorithm might overestimate the ratios for this protein, as for example visible for the 1K and Cyt fractions of the MCM7 trace, which was not the case for directLFQ (**Figure 4D**). Conversely, to investigate the cause of the discrepancy for the proteasome, we inspected all aligned precursor intensity traces that underlie the protein intensity estimation, which is only possible at the protein level in MaxLFQ. Note that the shapes of each intensity trace are untouched, therefore accurately reflecting the underlying data. This revealed that the protein intensity estimation is indeed consistent with the precursor data for both MCM7 as well as PSMB2 in directLFQ. For PSMB2 there are several datapoints supporting the variation in the shape of the protein intensity profile in the 6K fraction. Thus, despite the higher variance, the data clearly validate PSMB2 as a member of the proteasome complex, while at the same time indicating underlying technical or even biological reasons such as peptide modifications for the deviating shape.

## DISCUSSION

Here, we have introduced a simple yet effective algorithm for normalization and protein intensity estimation for DDA as well as DIA proteomics data. Its central concept is the shifting of peptide intensity traces and sample intensity traces on top of each other. In computer science terms, is of linear order O(n) where n can be the number of samples or the proteins. In practice this allows the quantification of extremely large sample sizes of hundreds of thousands. As with any other algorithm, there are hardware requirements that have to be fulfilled and in particular, directLFQ is currently memory limited, with a rough estimate of around 30GB of memory necessary for 10,000 samples. While not a practical problem now, in the future this could be alleviated by using standard out-of-memory computing approaches. The underlying code is openly available on GitHub under the Apache License facilitating improvements and contributions from the community. The package is easily accessible via Python Package Index (PyPI) as well as through one-click installers coupled to a graphical user interface. As the concepts underlying directLFQ are relatively straightforward, we hope for wider use of the algorithm within the community.

We show that directLFQ compares favorably in biological and technical benchmarks to the state-of-the-art implementations of MaxLFQ algorithms. Unlike these, directLFQ also provides normalized peptide or fragment-ion intensities which allows retracing the protein intensity estimation, enabling inspection of individual peptides.

The directLFQ approach effectively deals with quantification challenges such as differing sample loadings, differing ionization efficiencies between peptides as well as missing values. Nevertheless, as in the other current protein intensity estimation approaches, some challenges remain:

Firstly, directLFQ solves the ionization efficiency problem by using relative instead of absolute quantification. This is done by (implicitly) comparing the relative quantification of all samples. This stabilizes the protein intensity estimation, but also means that each individual sample can influence the protein intensity estimation of all other samples. In other words, actions like adding or removing samples may slightly alter the protein intensity estimations of all other samples. In practice, this means that all comparisons have to be done on a dataset that has been processed in the same directLFQ analysis.

Secondly, recall that the aim of directLFQ is to estimate protein intensities. By definition, this reduces multiple data points that could describe the protein to a single data point that is the “best guess” protein intensity. This reduces overall complexity and is useful for many tasks, such as globally analyzing protein behavior. However, this aspect of quantification only represents a subset of the tasks and algorithms required for comprehensive analysis of quantitative proteomics data. In particular, it is usually necessary to perform statistical analyses such as differential expression analysis in order to determine the regulation between biological conditions. For these, it can be more informative to retain the ion-level information (precursor or transition intensities) instead of working with protein intensity estimates ^31,35^. For this, resolving peptide level information more deeply is an outstanding challenge in quantitative proteomics. In particular, interferences, systematic biases and assessment of consistency from the basic ion-level to the protein or gene levels will still need to be better addressed, a topic that we are working on already.

The possibility to quantify increasingly large numbers of samples becomes more and more important with the advent of high-throughput proteomics approaches, such as measuring very large cohorts of patients. Additionally, the emerging field of single cell proteomics will drastically increase the number of cells (and therefore samples) that will be measured and quantified. With directLFQ we provide an approach that allows fast quantification for all such scenarios. Furthermore, directLFQ can already be used on other MS data types such as TMT and we see no reason why it should not be useful for RNA sequencing data, too.

## ABBREVIATIONS

|  |  |
| --- | --- |
| DDA | data-dependent acquisition |
| DIA | data-independent acquisition |
| CV | coefficient of variation |
| TMT | tandem mass tag |
| LFQ | label-free quantification |
| QC | quality control |
| DOM | dynamic organellar maps |
| PyPI | python package index |
| GUI | graphical user interface |
| API | application programming interface |

## DATA AVAILABILITY

All code to reproduce the results shown in this study is available under https://github.com/MannLabs/directlfq with downloaders for the underlying data provided there. The identifiers of the PRIDE repositories used to download the original data are provided in the respective Experimental Results sections. The MaxQuant results tables used for the organellar maps data can be downloaded under the PRIDE reviewer account provided with the submission.

## ACKNOWLEDGEMENTS

We thank our collages in department of Proteomics and Signal Transduction and the Clinical Proteomics group at the NNF Center for Protein Research for help and fruitful discussions, especially, Patricia Skowronek, Marvin Thielert, Maximilian Strauss and Jakob Bader.

This study was supported by The Max-Planck Society for the Advancement of Science. C.A., S.W. and A.M. acknowledge funding from the Bavarian State Ministry of Health and Care as part of “DigiMed Bayern” (grant No: DMB-1805-0001).

## AUTHOR CONTRIBUTIONS

C.A. and M.M. conceptualized the directLFQ approach. C.A. designed, implemented and evaluated the algorithm. J.P.S. contributed to the analysis of the organellar maps data. A.C.M. contributed to the execution time analyses. S.W. provided important infrastructure code. C.A. and M.M. wrote the paper with input from all authors.

## SUPPLEMENTARY FIGURES

**Supplementary Figure 1:**
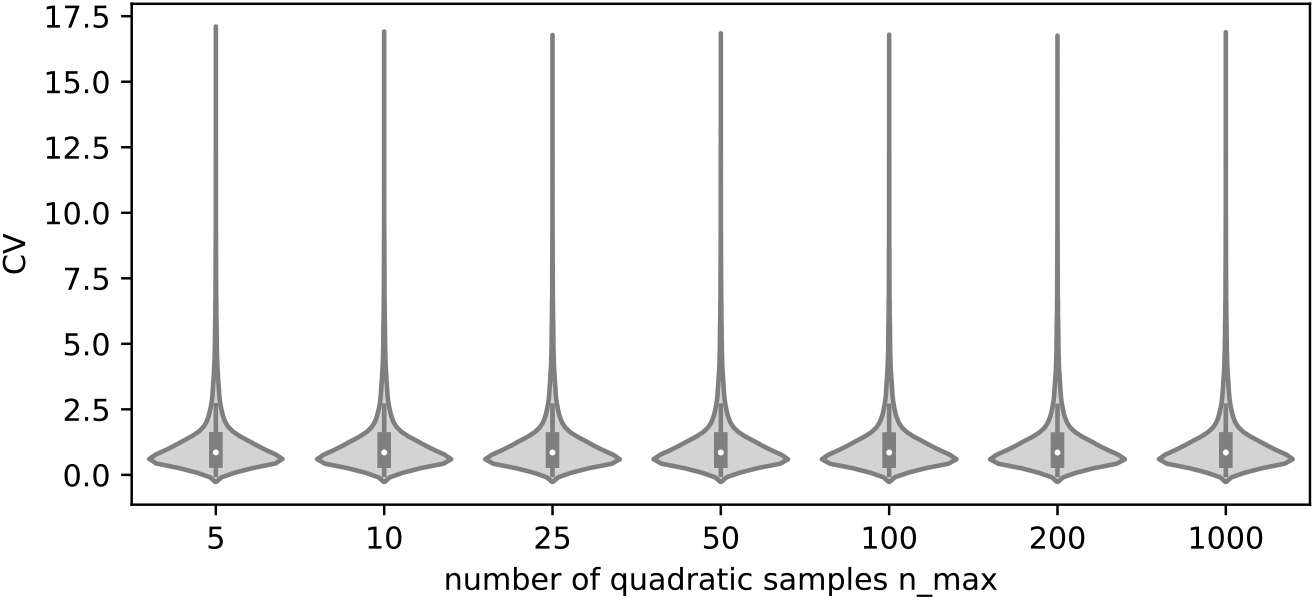
CVs over precursors of the yeast interactome datasets, for different number of samples *n_max_* used for creating an average trace as described in the main text. No visible change depening on *n_max_* can be seen in these distributions, indicating that a high *n_max_* is not necessary for improved performance.

**Supplementary Figure 2:**
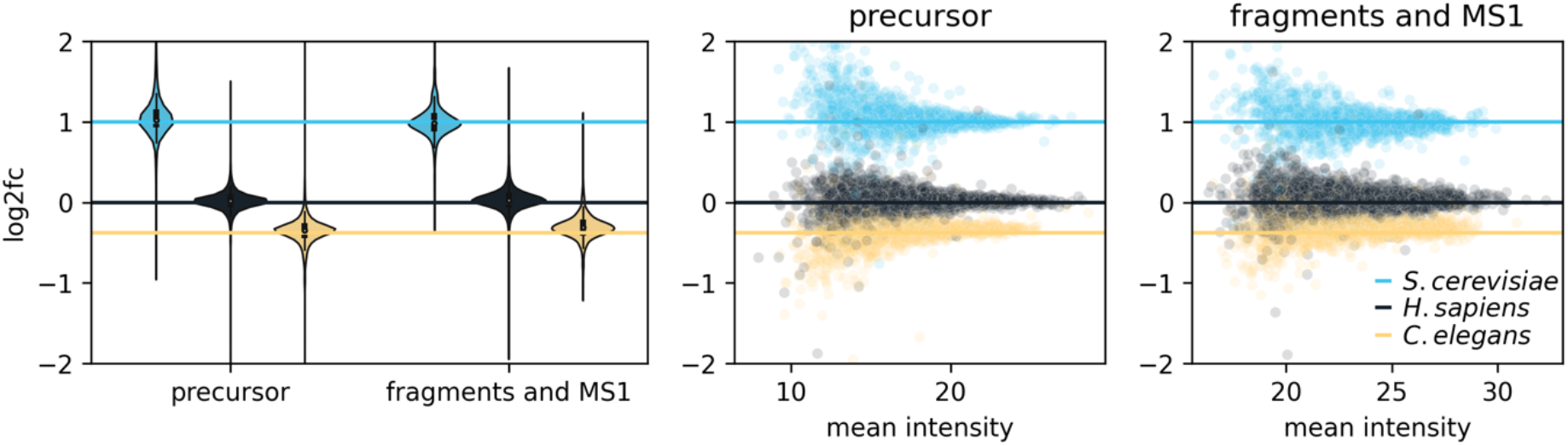
Comparing protein quantification in DIA based on precursor-as well as fragment and MS1 level data. The underlying dataset is the mixed species experiment of **Figure 3C** and described in the main text. In the case of precursor-level data, the directLFQ intensity traces are derived from the “FG.Quantity” column in the Spectronaut report, which gives one intensity per charged and potentially modified peptide sequence. The fragment and MS1 level data assigns one intensity trace to each individual fragment ion (“F.PeakArea”) and to each MS1 isotope intensity (“FG.MS1IsotopeIntensities (Measured)”). Overall both results result in good quantification, with proteins meeting the expected fold changes, but the fragment seems to be more consistent. For this reason, we chose the fragment approach for DIA data analysis in this study. The user does however have the option to analyze based on precursor information.

**Supplementary Figure 3:**
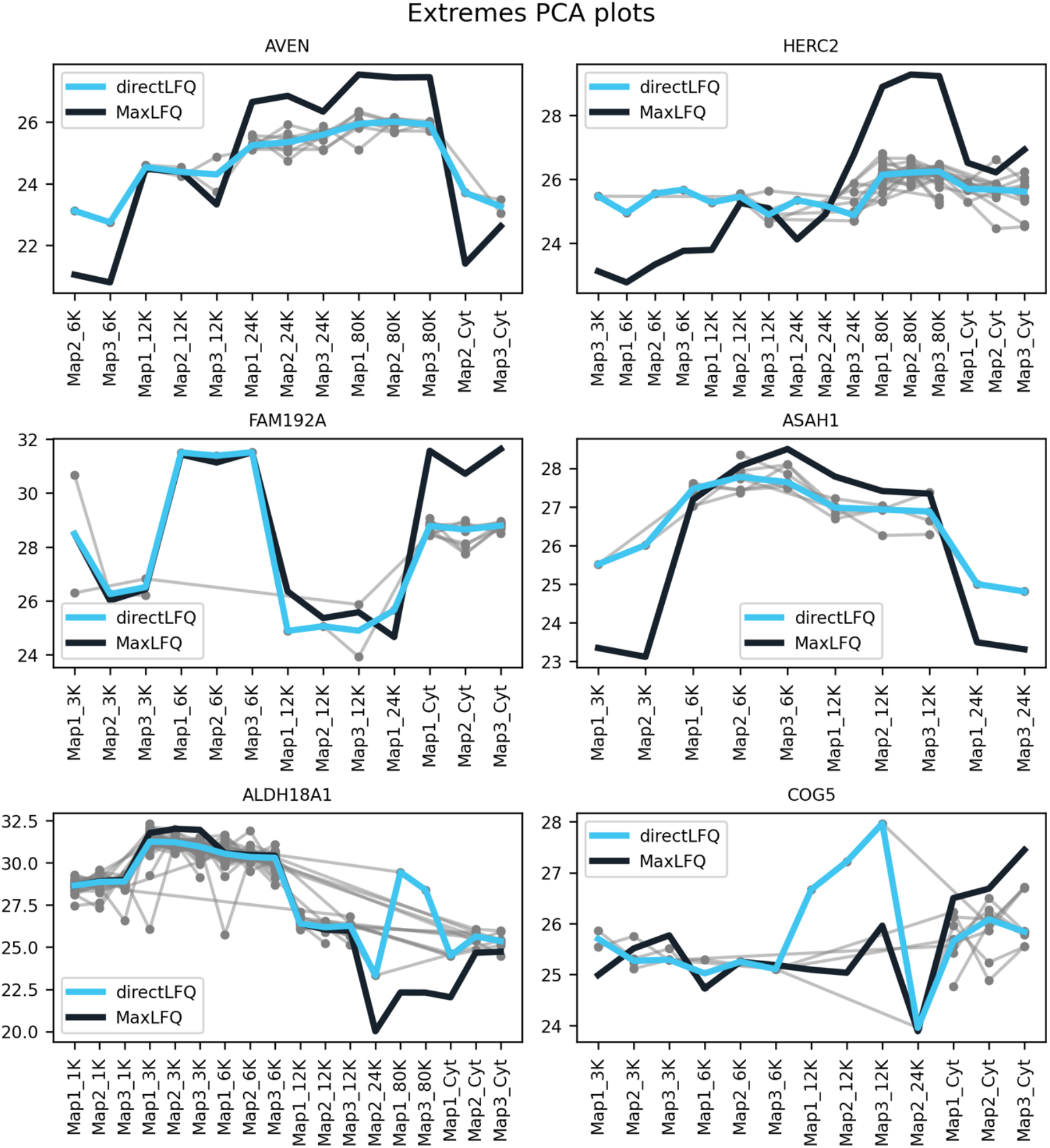
Visualizing estimated protein intensity traces for MaxLFQ and directLFQ for different proteins (see main text for a more detailed explanation of the ion intensity plot). The proteins selected were on the edges of the PCA plots displayed in **Figure 4A&B**. We deemed such outlying proteins to be good example cases to see the differences between the directLFQ and the MaxLFQ approach. We see directLFQ (by definition) consistent with the underlying normalized ion data. The MaxLFQ traces are harder to fit towards the normalized ion traces shown here and seem to display stronger variations.

